# Point contact-restricted cAMP signaling control ephrin-A5-induced axon repulsion

**DOI:** 10.1101/2024.04.20.589861

**Authors:** J Bécret, C Michaud, A Assali, NAL Chenais, I Kankadze, C Gomez-Bravo, F Roche, S Couvet, C Fassier, X Nicol

## Abstract

Signal transduction downstream of axon guidance molecules is essential to steer developing axons. Second messengers including cAMP are key molecules shared by a multitude of signaling pathways and are required for a wide range of cellular processes including axon pathfinding. Yet, how these signaling molecules achieve specificity for each of their downstream pathways remains elusive. Subcellular compartmentation emerged as a flexible strategy to reach such a specificity. Here, we show that point contact-restricted cAMP signals control ephrin-A5-evoked axon repulsion in vitro by modulating Focal Adhesion Kinase phosphorylation and the assembly and disassembly rate of point contacts. Consistently, preventing point contact-specific cAMP signals, in developing retinal ganglion cells in vivo alters the refinement of their terminal axonal arbor in the brain. Altogether, our study identifies point contacts as a compartment containing a local cAMP signal required for ephrin-A5-dependent axon guidance and highlights the crucial role of such subcellularly restricted second messenger signals in the wiring of neuronal circuits.

## Introduction

cAMP is a key signaling molecule influencing a wide range of cellular events including the wiring of developing neuronal circuits. Nevertheless, it controls each of its downstream pathways with high specificity. The subcellular compartmentation of this second messenger contributes to the selective activation of its downstream effectors, although cAMP freely diffuses in aqueous buffers. The combination of local cAMP synthesis by adenylyl cyclases combined with hydrolysis by phosphodiesterases contributes to its subcellular compartmentation ^1–3^. In developing neurons, a few cellular compartments with distinct cAMP signals have been identified. These cellular domains include for instance the presumptive axon and the future dendrites that exhibit distinct and interdependent cAMP signals during axonogenesis ^4^. During axon pathfinding, cAMP signaling is also highly compartmentalized. While developing axons grow towards their targets, repulsive and attractive cues orient their growth. Interpretation of these extracellular cues relies on second messenger signals restricted to specific cellular domains. For instance, cAMP signaling in lipid rafts is required downstream of the axon repellent ephrin-A5 to refine the terminal arbors of retinal axons ^5^. In contrast, the axonal repulsion elicited by another guidance molecule, Slit1, is required for retinal axon guidance at the chiasm and relies on cAMP modulation in a microdomain of the plasma membrane that does not overlap with lipid rafts ^6^. However, the molecular and cellular events controlled by each local cAMP signaling in navigating axons have not been identified. Additionally, while cAMP signals confined in at least two distinct subcellular compartments drive specific axon behaviors, it is still unclear whether modulations of cAMP signals restricted to other cellular domains contribute to axon guidance.

The regulation of cellular adhesion is a central mechanism that controls axon pathfinding ^7^. Among the modalities that influence the interaction of axonal growth cones with their substrate, integrin-dependent adhesion is critical for axon guidance. When bound to their ligand in the extracellular matrix, integrins induce the assembly of macromolecular complexes called focal adhesions in non-neuronal cells. Focal adhesions connect the extracellular matrix and integrins to the actin cytoskeleton. In axons, the integrin-associated macromolecular complexes are termed point contacts (PCs). PCs have a smaller size, less elongated shape and shorter lifetime than focal adhesions, but share many molecular components with the early stages of these molecular complexes ^8^. Focal Adhesion Kinase (FAK) is a master regulator of both focal adhesions and PC dynamics and functions. In developing growth cones and in non-neuronal cells, FAK is critical for the cellular response to axon guidance molecules including ephrin-As ^9,10^, Netrin-1 ^11,12^ and Semaphorin-3A ^13^. Notably, this kinase controls the pathfinding of retinal axons as well as the topographic positioning of their terminal arbor in non-mammalian vertebrate models, a process that requires ephrin-A signaling ^10,14^.

Interestingly, cAMP signaling indirectly influences FAK phosphorylation ^15^. For instance, the cAMP-dependent protein kinase PKA is involved in cannabinoid-induced FAK phosphorylation in hippocampal neurons ^16^. This prompted us to investigate whether subcellularly-restricted cAMP signals regulate cellular adhesion by controlling the remodeling of PCs in growth cones exposed to axon guidance molecules. We here identified PCs as a subcellular domain in which cAMP modulation is required for axon guidance. We show that PC-specific cAMP signaling controls ephrin-A5-induced retinal axon retraction in vitro, through the modulation of FAK phosphorylation and the regulation of PC turn-over. We also demonstrate that this local cAMP signaling is required for the development of retinal axons in vivo by shaping their terminal arbor.

## Results

### PC-restricted cAMP signaling is required for ephrin-A5-induced axon retraction

To evaluate whether cAMP modulation in PCs influences retinal ganglion cell (RGC) axon pathfinding, we developed a PC-restricted variant of cAMP Sponge, a genetically-encoded cAMP scavenger. cAMP Sponge expression acts as a cAMP buffer and prevents the modulation of cAMP downstream pathways. Its expression can be restricted to chosen subcellular compartments by fusing it to appropriate targeting sequences ^5,6,17^. Here, cAMP sponge was fused to paxillin (pax-cAMP Sponge, **Figure 1a**), a central component of both focal adhesions and PCs. This strategy has previously been used to target exogenous proteins to focal adhesions in other cell types ^18^. Pax-cAMP Sponge also includes a mCherry tag to identify the neurons expressing this PC-targeted cAMP buffer. The targeting of pax-cAMP Sponge to PCs was verified by evaluating the colocalization of pax-cAMP Sponge with the full length paxillin fused to GFP, a previously reported PC marker (pax-GFP) ^19^. In the growth cone of RGC developing axons, pax-cAMP Sponge and pax-mRFP colocalize to a similar extent with pax-GFP, confirming the targeting of pax-cAMP Sponge to PCs (**Supplementary Figure 1**). Pax-cAMP Sponge was expressed in embryonic RGCs in vitro to investigate the consequences of altered cAMP signaling in PCs on the dynamic behavior of developing RGC axons exposed to ephrin-A5. The morphological changes of control axons exposed to this guidance cue were characterized by a prompt lamellipodium loss (growth cone collapse) followed by the elimination of filopodia and the retraction of the axon (**Figure 1b**). Preventing changes in cAMP signaling in PCs by expressing pax-cAMP Sponge did not affect the collapse of the growth cone but reduced axon retraction (**Figure 1b**), demonstrating the influence of PC-restricted cAMP signals on the ephrin-A5-induced backward movement of the growth cone. This phenotype is in line with the previously described impact of non-compartmented cAMP manipulations on ephrin-A-dependent axon pathfinding ^20,21^.

**Figure 1.**
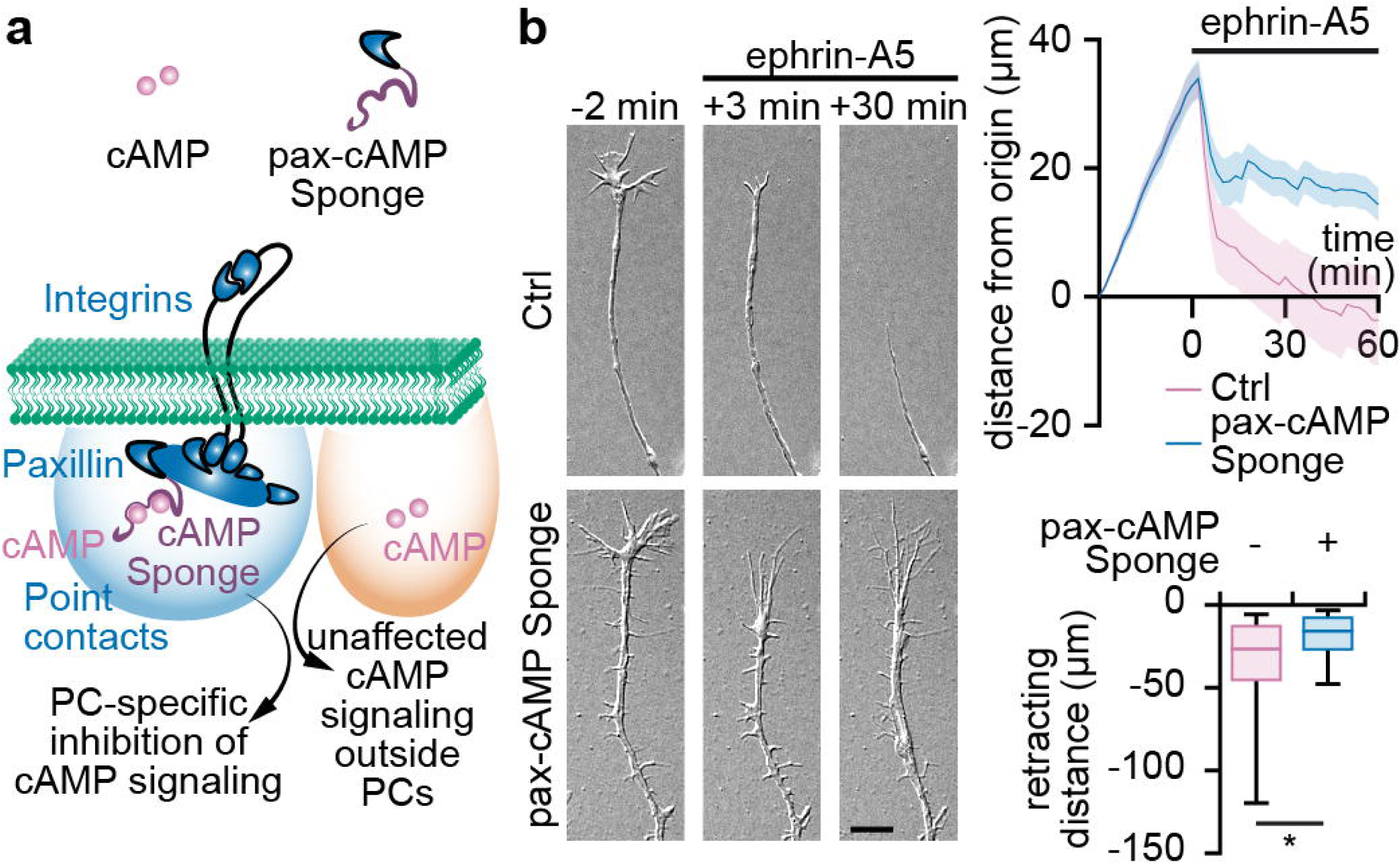
PC-restricted cAMP signaling is required for ephrin-A5-induced axon retraction. **(a)** Approach for PC-restricted manipulation of cAMP signaling. The genetically-encoded cAMP scavenger cAMP Sponge was fused to paxillin (pax-cAMP Sponge) to restrict its expression to PCs and locally prevent modulations of cAMP signaling. **(b)** Still images of live DIC imaging of RGC axons expressing pax-cAMP Sponge or pax-RFP (Ctrl). Pax-cAMP Sponge impairs ephrin-A5-evoked axon retraction. **(b)** Traces: mean ± s.e.m. Box-and-whisker plot elements: median, upper and lower quartiles, 10^th^ and 90^th^ percentiles. * P<0.05, two-tailed Mann-Whitney test. Scale bar, **(b)** 10 µm.

### PC-restricted cAMP signaling is required for long term reduction of PC density

To identify the cellular event controlled by PC-restricted cAMP signaling in axons exposed to guidance molecules, we investigated the changes in PC density induced by ephrin-A5. Axons were stained for a major component of PCs, focal adhesion kinase (FAK), 5 min and 1 hour after the onset of ephrin-A5 exposure to evaluate the short- and long-term impact of PC-restricted cAMP manipulation on PC density. Axons were imaged using TIRF microscopy to ensure that the analysis was restricted to the FAK population that is adjacent to the plasma membrane, i.e. engaged in cellular adhesion. This approach excludes recycling or trafficking FAK proteins. The density of FAK-positive puncta was reduced in control (pax-RFP-expressing) axons exposed 5 minutes to ephrin-A5. No significant difference was detected in ephrin-A5-treated axons expressing pax-cAMP Sponge as compared to their pax-RFP-expressing controls (**Figure 2a**). After 1 hour of ephrin-A5 exposure, the density of FAK-expressing PCs was reduced in pax-RFP-expressing growth cones, demonstrating a continuous reduction of the density of PCs over time (**Figure 2b,c**). In contrast, the density of FAK-positive PCs was higher in pax-cAMP Sponge expressing axons as compared to control axons (**Figure 2c,d**). The higher PC density observed in pax-cAMP Sponge-expressing axons 1 hour after ephrin-A5 exposure was confirmed with another PC marker, paxillin. Similarly, to FAK-positive PCs, the density of paxillin-expressing PCs was reduced 1 hour after the onset of ephrin-A5 treatment in controls. In contrast, this reduction in the density of paxillin-positive puncta was not detected in growth cones expressing pax-cAMP Sponge (**Figure 2d**).

**Figure 2.**
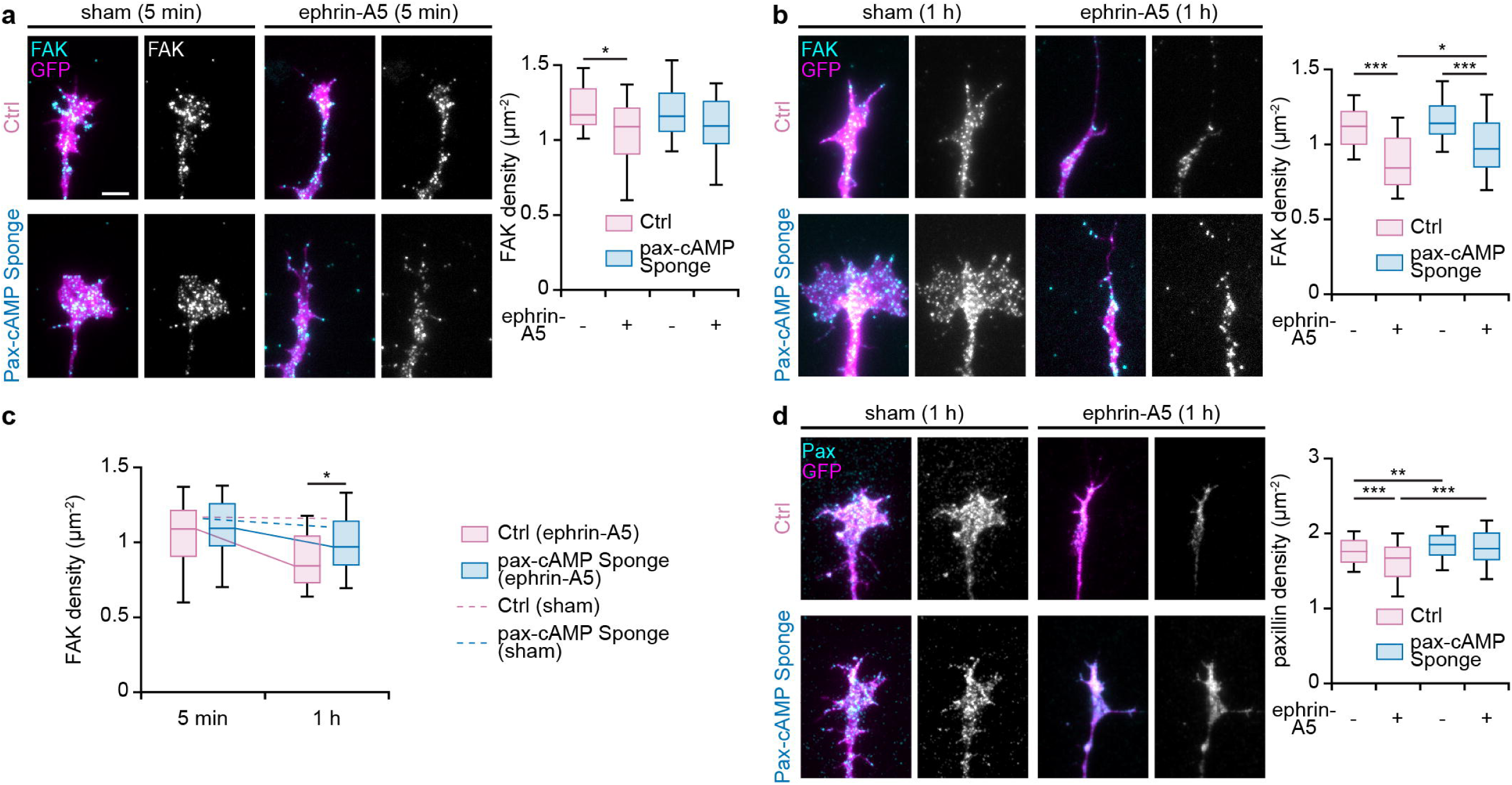
PC-restricted cAMP signaling is required for the ephrin-A5-induced reduction in the density of FAK-positive PCs. **(a)** Retinal axons expressing GFP together with either pax-RFP (Ctrl) or pax-cAMP Sponge were exposed to ephrin-A5 or PBS as a sham control, immunolabeled for FAK and imaged using TIRF microscopy. Ephrin-A5 induces a reduction in the density of FAK 5 min after the stimulation of control growth cones. This reduction was not detected in pax-cAMP Sponge-expressing axons. However, no statistically significant difference was detected between control and pax-cAMP Sponge- expressing axons. **(b)** Long term exposure to ephrin-A5 (1 h) leads to a further reduction in FAK- containing PCs in control axons. This decrease of FAK-positive puncta is reduced in pax-cAMP Sponge-expressing growth cones. **(c)** Summary of the changes in FAK-positive PCs over time. **(d)** After 1 hour of ephrin-A5 exposure, the density of paxillin-positive PCs is reduced in pax-RFP- expressing axons, but not in axons electroporated with pax-cAMP Sponge. Box-and-whisker plot elements: median, upper and lower quartiles, 10^th^ and 90^th^ percentiles. * P<0.05; ** P<0.01; *** P<0.001; Kruskal-Wallis test followed by Dunn’s post-hoc test. Scale bar, 5 µm.

Overall, these observations demonstrate that PC-restricted cAMP signaling is critical for the ephrin-A5-evoked reduction of PC density.

### Influence of PC-restricted cAMP signaling on PC turnover

The dynamics of PCs critically influences their density. Indeed, the latter is highly dependent on the assembly and disassembly rate of these adhesion-dedicated structure and is influenced by their lifetime. To evaluate whether ephrin-A5 influences the dynamics of PCs in RGC axons and whether PC-restricted cAMP signaling is involved in this process, we electroporated RGCs with a plasmid encoding paxillin fused to GFP (pax-GFP). This approach enables to monitor the turnover of PCs in living retinal growth cones while being exposed to ephrin-A5 in vitro. Axons were monitored using TIRF imaging of pax-GFP to ensure tracking only paxillin contributing to cellular adhesion. In control axons, the density of pax-GFP assemblies is gradually reduced 5 and 10 minutes after the onset of ephrin-A5 stimulation (**Figure 3a,b**). The reduction in the density of pax-GFP puncta is delayed by the expression of pax-cAMP Sponge and the number of paxillin-containing molecular assemblies is higher than in control axons 10 minutes after ephrin-A5 exposure (**Figure 3a,b**). These observations match the overall conclusions made using FAK antibodies (**Figure 2**).

**Figure 3.**
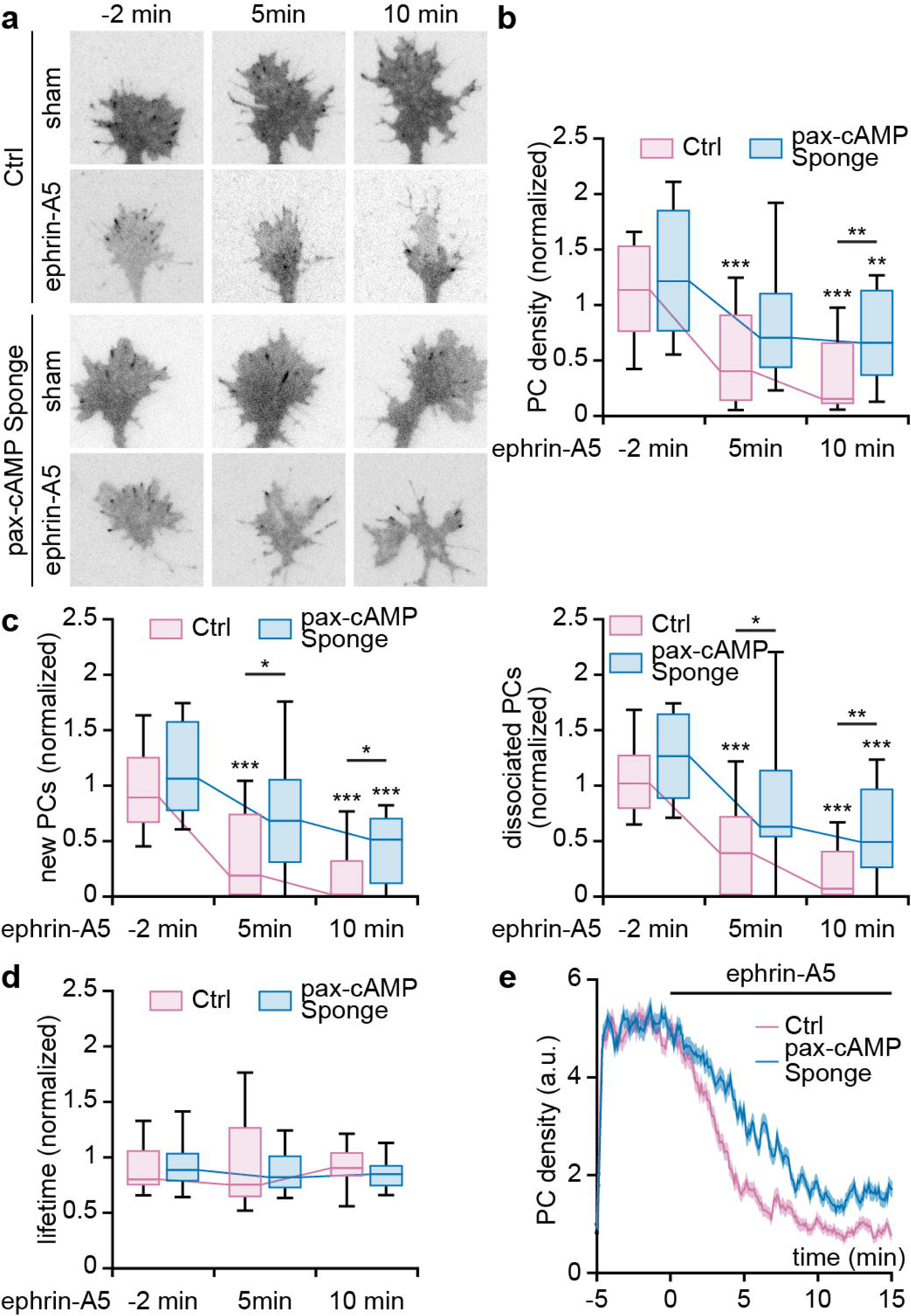
cAMP signaling in PCs is required for the ephrin-A5-induced change of PC dynamics. **(a)** Living pax-RFP- and pax-cAMP Sponge-expressing axons were imaged using TIRF microscopy while being exposed to ephrin-A5. Axons co-expressed pax-GFP and this reporter was imaged in the same growth cone before, 5 minutes and 10 minutes after the stimulation. **(b)** The density of pax-GFP-positive PCs is reduced over time by ephrin-A5. This reduction is attenuated when pax-cAMP Sponge is expressed. **(c)** Similarly, the appearance and dissociation rate of PCs containing pax-GFP is lowered by ephrin-A5, and this trend is reduced by pax-cAMP Sponge expression. **(d)** In contrast, neither ephrin-A5 nor pax-cAMP Sponge influences the lifetime of pax-GFP-positive PCs. **(e)** Model mimicking the evolution of lifetime, appearance rate and dissociation frequency of PCs. The model uses the experimental measurements of these parameters and highlights that a lower reduction of both the appearance and dissociation rate leads to a higher density of PCs. 100 simulations rounds are average. Box-and-whisker plot elements: median, upper and lower quartiles, 10^th^ and 90^th^ percentiles. * P<0.05; ** P<0.01; *** P<0.001; paired comparisons: Friedman test, unpaired comparisons: Mann-Whitney. Scale bar, 10 µm. Shadows around traces represent s.e.m.

To evaluate if the changes in PC turnover can explain the lower reduction in PC density associated with cAMP signaling buffering in this subcellular compartment, the frequency of assembly and disassembly of pax-GFP-positive puncta was quantified using live TIRF imaging. Ephrin-A5 induces a drastic and sustained drop of both PC assembly and disassembly rate 5 minutes after stimulation (**Figure 3c**). These reductions of the assembly and disassembly frequencies were less pronounced in pax-cAMP Sponge-expressing axons (**Figure 3c**). In contrast, the lifetime of the monitored pax-GFP-positive PCs was neither affected by ephrin-A5 exposure nor by pax-cAMP Sponge expression (**Figure 3d**).

Explaining a reduction in the density of PCs by similar reductions in their assembly and disassembly rates without affecting their lifetime may sound counterintuitive. A simple mathematical simulation was developed to evaluate if the measured changes in PC dynamics are sufficient to explain the ephrin-A5-induced changes in pax-GFP puncta density and the impact of pax-cAMP Sponge expression. The model runs PC assembly and disassembly cycles with the lifetime, assembly and disassembly rates adjusted to the measures obtained from live TIRF imaging. Using these parameters, the simulation mimics the evolution of PC density in control and pax-cAMP Sponge-expressing axons. It also confirmed that reducing the rate of assembly and disassembly without affecting their ratio (close to 1) and without changing the lifetime of PCs leads to a decrease in the PC density (**Figure 3e**). It further provides an explanation linking the reduced decrease of PC assembly and disassembly rates in pax-cAMP Sponge-expressing axons to the higher PC density as compared to control axons exposed to ephrin-A5.

Overall, these results demonstrate that PC-restricted cAMP signaling is required for the ephrin-A5-mediated control of PC turnover and the subsequent reduction in the density of these molecular complex that is associated with axonal retraction.

### PC-restricted cAMP signaling specifically induces FAK phosphorylation on tyrosine 925

To identify the molecular events underlying PC remodeling by PC-restricted cAMP signaling, we focused on FAK phosphorylation. FAK carries a large set of phosphorylation sites including some that have been involved in axon pathfinding. While FAK autophosphorylation on tyrosine 397 influences both axon attraction and repulsion ^10–13^, the phosphorylation of tyrosines 861 and 925 selectively favors axon attraction and repulsion, respectively ^11–13,22,23^. To determine whether these phosphorylations are influenced by ephrin-A5 and by PC-restricted cAMP signaling, control or pax- cAMP Sponge-expressing retinal axons were exposed to ephrin-A5 during 5 minutes or 1 hour and immunostained with antibodies specifically raised against FAK phospho-tyrosine 925 (^p925^FAK), 861 (^p861^FAK) or 397 (^p397^FAK). In control axons, ephrin-A5 induces a decrease in the density of PCs containing ^p925^FAK, ^p861^FAK or ^p397^FAK from 5 minutes post-treatment (**Figure 4a, Supplementary Figure 2a,d**). These density reductions were still detected 1 hour after ephrin-A5 exposure and were more pronounced for ^p861^FAK and ^p397^FAK (**Figure 4b,c, Supplementary Figure 2b,c,e,f**). The density of ^p925^FAK-, ^p861^FAK- and ^p397^FAK-containing PCs was similar in controls and pax-cAMP Sponge-expressing growth cones after 5 min of ephrin-A5 exposure (**Figure 4a, Supplementary Figure 2a,d**). In contrast, cAMP buffering in PCs selectively amplifies the reduction in the density of ^p925^FAK-positive but not of ^p861^FAK- and ^p397^FAK-labelled PCs 1 hour after the onset of the treatment (**Supplementary Figure 2b,c,e,f**).

**Figure 4.**
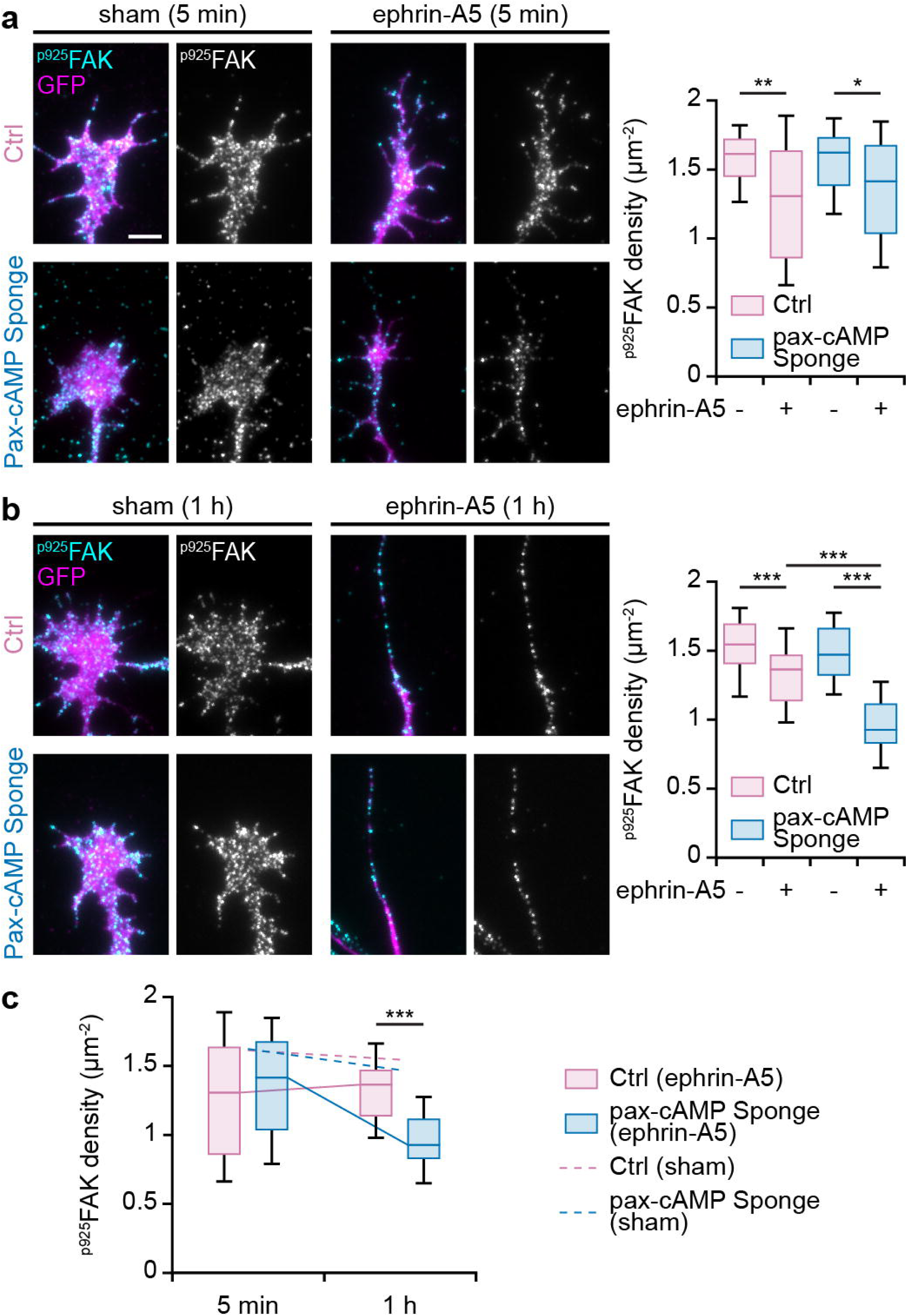
PC-restricted cAMP signaling is required for the phosphorylation of FAK on tyrosine 925. Retinal axons expressing GFP together with either pax-RFP or pax-cAMP Sponge were exposed to ephrin-A5 or PBS as a sham control, immunolabeled for the phosphorylated tyrosine 925 of FAK and imaged using TIRF microscopy. **(a)** Ephrin-A5 induces a reduction in the density of ^p925^FAK 5 min after the stimulation, in both control and pax-cAMP Sponge-expressing growth cones. **(b)** Long term ephrin-A5 exposure (1 h) leads to a further reduction in ^p925^FAK-containing PCs in pax-cAMP Sponge-expressing axons, in contrast to pax-RFP-electroporated axons. **(c)** Summary of the changes in ^p925^FAK-positive PCs over time. Box-and-whisker plot elements: median, upper and lower quartiles, 10^th^ and 90^th^ percentiles. * P<0.05; ** P<0.01; *** P<0.001; Kruskal-Wallis test followed by Dunn’s post-hoc test. Scale bar, 5 µm.

Overall, our results demonstrate that compartmented cAMP signaling in PCs influences the phosphorylation of FAK on the tyrosine 925 during ephrin-A5-induced axon retraction. They further suggest that PCs containing ^p925^FAK before ephrin-A5 exposure may disassemble within 5 minutes after ephrin-A5 stimulation in a cAMP-independent mechanism thus explaining the reduction in ^p925^FAK density both in control and pax-cAMP Sponge-expressing growth cones. Further tyrosine 925 phosphorylation may be induced by sustained ephrin-A5 exposure through a mechanism requiring PC-restricted cAMP signaling, explaining the selective amplified reduction in ^p925^FAK when cAMP is buffered in PCs.

### Preventing the phosphorylation of FAK at Y925 mimics the absence of cAMP signaling in PCs

To assess whether the control of FAK phosphorylation on tyrosine 925 by PC-restricted cAMP modulation is linked to ephrin-A5-induced axon retraction, a dominant-negative form of FAK that cannot be phosphorylated on the tyrosine 925 (^Y925F^FAK phosphoresistant variant) was electroporated in RGCs. FAK phosphoresistant mutants for tyrosine 397 or 861 (^Y397F^FAK and ^Y861F^FAK, respectively) were used to evaluate the impact of these cAMP signaling-independent phosphorylation sites ^24^. Retinal axons expressing each phosphoresistant variant were exposed to ephrin-A5 for 1 hour and the percentage of collapsed axons as well as the axon retraction length were measured in each experimental condition. The latter was evaluated by measuring the longest remaining filopodia in collapsed growth cones, a measurement that has been previously identified as a proxy for axon retraction ^21^. None of the phosphoresistant FAK variants impact ephrin-A5-evoked growth cone collapse (**Figure 5**). In contrast, both ^Y397F^FAK and ^Y925F^FAK phosphomutants alter the axonal retraction elicited by this repulsive cue unlike ^Y861F^FAK or the overexpression of wild-type FAK (**Figure 5 and Supplementary Figure 2**). The retraction defect associated with ^Y397F^FAK and ^Y925F^FAK is illustrated by the reduced length of the retraction process, i.e. the remaining filopodia that is distal to the retraction bulb (**Figure 5**). These observations are consistent with previous reports linking ^p925^FAK and ^p397^FAK with axon repulsion events ^10–13,22,23^. Altogether our results demonstrate that ephrin-A5-induced axon retraction requires FAK phosphorylation at tyrosine 925 through a mechanism dependent on PC-restricted cAMP signaling. While also required for this cellular process, FAK phosphorylation at tyrosine 397 occurs independently of this local cAMP signal (**Supplementary Figure 2**).

**Figure 5.**
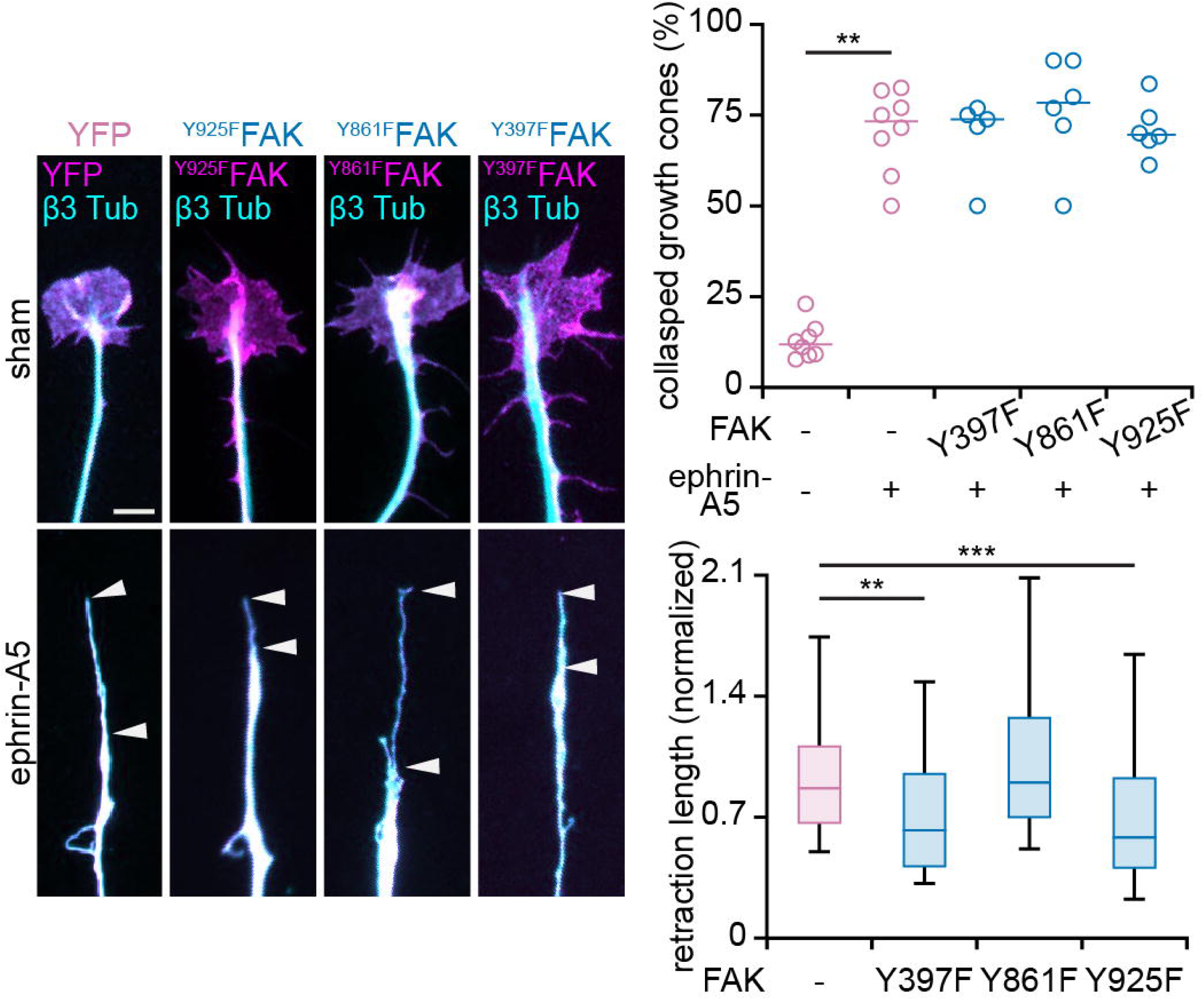
Preventing FAK phosphorylation on tyrosine 925 or 397 reduces ephrin-A5-induced axon retraction. Axons expressing phosphoresistant forms of FAK (^Y925F^FAK, ^Y397F^FAK, ^Y861F^FAK) were exposed to ephrin-A5 for 1 hour. Neither ^Y925F^FAK nor ^Y397F^FAK nor ^Y861F^FAK influence the number of collapsed growth cones. ^Y925F^FAK- and ^Y397F^FAK- but not ^Y861F^FAK expression led to the shortening of the retraction process (between arrowheads), indicating that the retraction of these axons is reduced by these phosphoresistant forms of FAK. Top graph; median and individual values. Bottom graph; Box- and-whisker plot elements: median, upper and lower quartiles, 10^th^ and 90^th^ percentiles. ** P<0.01; *** P<0.001; Kruskal-Wallis test followed by Dunn’s post-hoc test. Scale bar, 5 µm.

### PC-restricted cAMP signaling drives distinct molecular events from lipid raft-specific cAMP pathway

Lipid raft-restricted cAMP signaling has been previously associated with ephrin-A5-induced retinal axon retraction ^5^. The phenotypic similarities between buffering cAMP in lipid rafts and PCs in ephrin-A5-exposed axons raised the question of whether these two cellular domains are features of the same compartment containing a single local cAMP signal. To test this hypothesis, we assessed the requirement of lipid raft-restricted cAMP signaling for the changes in ^p925^FAK phosphorylation induced by ephrin-A5 and dependent on PC-restricted cAMP signaling. cAMP signal was specifically prevented in lipid rafts of retinal axons using Lyn-cAMP Sponge, a cAMP scavenger targeted to this compartment ^5^. The density of PCs containing Y925-phosphorylated FAK was analyzed in retinal growth cones after 5 min and 1 hour of ephrin-A5 exposure. Buffering cAMP signaling in lipid raft does not affect the density of ^p925^FAK-positive PCs. Indeed, Lyn-cAMP Sponge-expressing axons exhibit similar reductions in the density of ^p925^FAK-positive PCs as control axons once exposed to ephrin-A5 (**Figure 6a-c**). This observation contrasts with the enhanced reduction in ^p925^FAK density observed in axons expressing pax-cAMP Sponge after one hour of ephrin-A5 exposure (**Figure 4b,c**), suggesting that lipid rafts and PCs are sites of distinct ephrin-A5-induced cAMP-signaling required for specific molecular events driving axonal repulsion.

**Figure 6:**
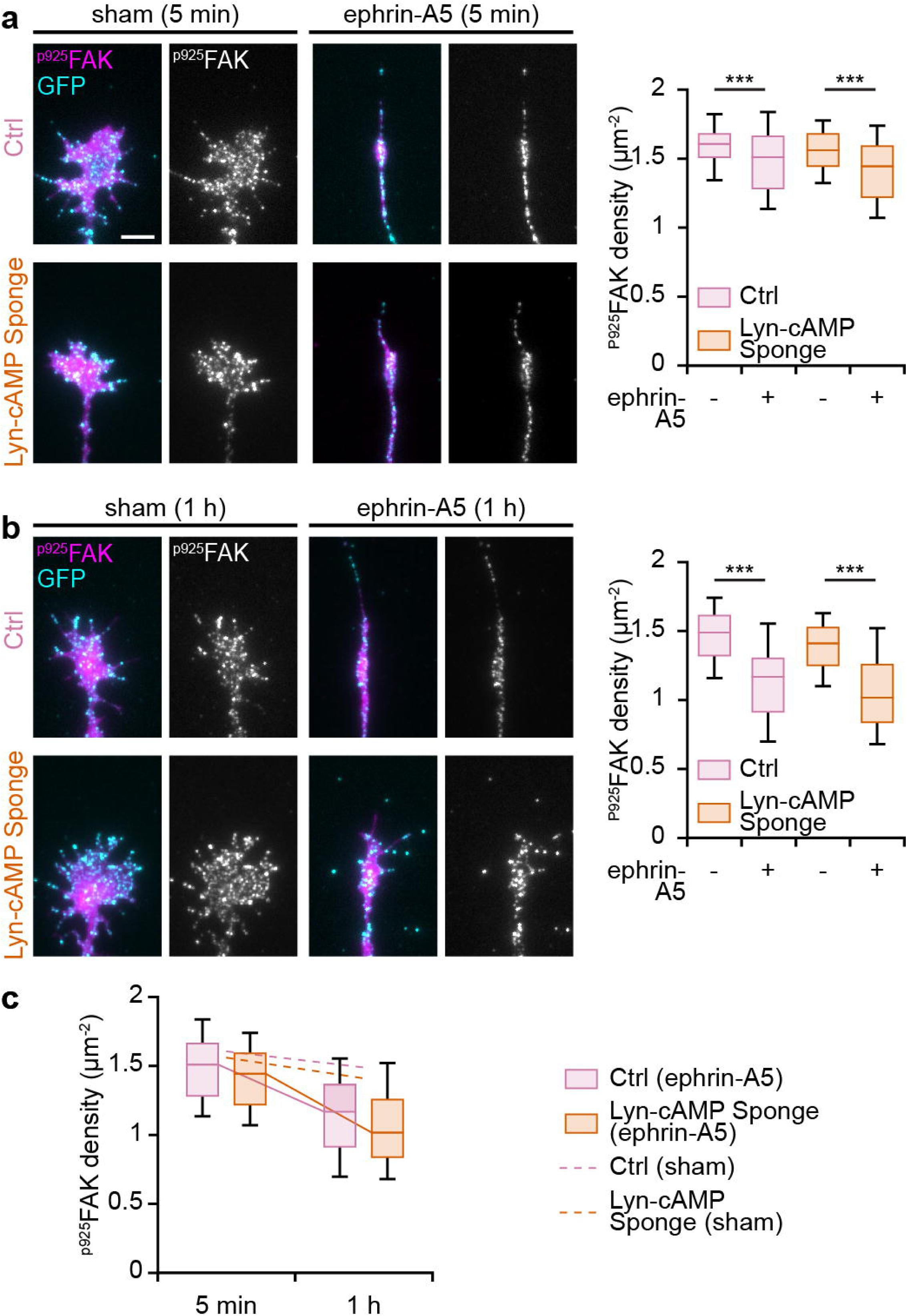
Lipid raft-restricted cAMP signaling does not influence ephrin-A5-induced changes in FAK phosphorylation on tyrosine 925. TIRF images of retinal axons expressing a lipid raft-targeted cAMP scavenger (Lyn-cAMP Sponge) or its mutated variant unable to bind cAMP (Lyn-mut cAMP Sponge) together with the GFP exposed to ephrin-A5 or PBS (sham control) and immunolabelled for phosphorylated tyrosine 925 of FAK. **(a)** Ephrin-A5 induces a reduction in the density of ^p925^FAK 5 min after the stimulation in both Lyn- cAMP Sponge-expressing and control growth cones. **(b)** Long term exposure to ephrin-A5 (1 h) leads to a reduction in ^p925^FAK-containing PCs that is similar in Lyn-cAMP Sponge- and Lyn-mut cAMP Sponge-expressing growth cones. **(c)** Summary of the changes in ^p925^FAK-positive PCs over time. Box-and-whisker plot elements: median, upper and lower quartiles, 10^th^ and 90^th^ percentiles. ** P<0.01; *** P<0.001; Kruskal-Wallis test followed by Dunn’s post-hoc test. Scale bar, 5 µm.

### PC-restricted cAMP signaling shapes the terminal arbor of retinal axons in vivo

To evaluate the impact of PC-restricted altered cAMP signaling on axon guidance in vivo, we electroporated pax-cAMP Sponge in utero in one retina of E14.5 mouse embryos and evaluated the development of RGC axon terminal arbors in the superior colliculus (SC) of P10 pups, a process that requires ephrin-A signaling. For individual axons, the end of each terminal branch was positioned in the SC and the volume they cover was assessed by computing the 3D convex hull of the identified branch ends. Pax-cAMP Sponge-expressing axons exhibit a similar number of branches as their controls. However, the end of their axonal branches were spread over a larger volume, demonstrating that Pax-cAMP Sponge-expressing retinal arbors are exuberant (**Figure 7a,b)**. This observation is in line with the reduced retraction of retinal axons exposed to ephrin-A5 in vitro.

**Figure 7.**
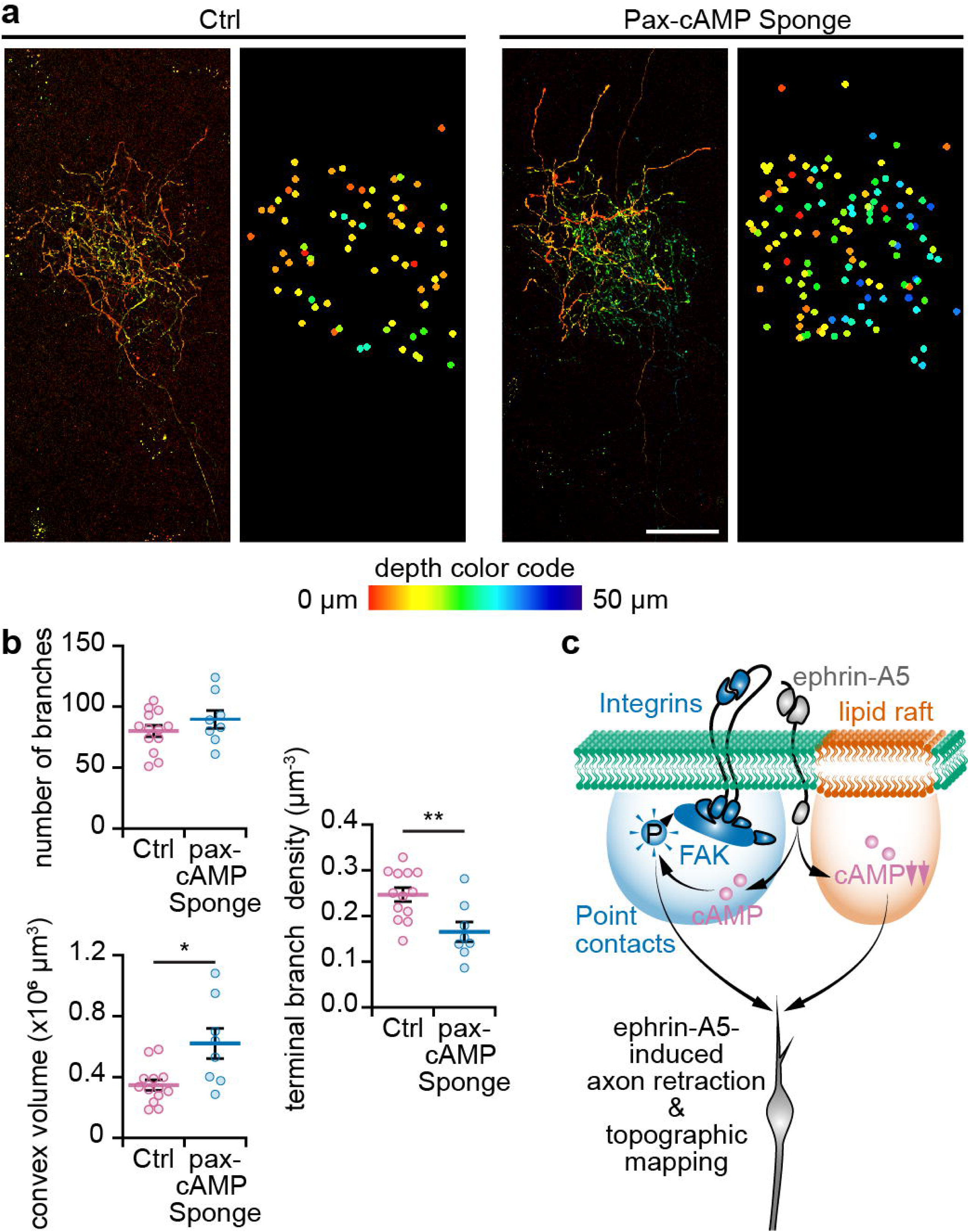
PC-specific cAMP signaling shapes the terminal arbor of retinal axons. **(a)** Confocal reconstructions of individual Ctrl (pax-RFP) and pax-cAMP Sponge-expressing axonal arbors in the SC. The depth is color-coded from red to blue. Dots in the right image highlight the position of the end of each terminal branch with the same depth color-code. **(b)** The terminal arbor of Pax-cAMP Sponge-expressing axons are spread over a larger volume, although the overall number of terminal branches is similar to control, thus leading to a reduced density of terminal branches. **(c)** Model of axonal cAMP signaling downstream of ephrin-A5. PC- and lipid raft-restricted cAMP signals contribute to axon retraction in vitro and to shaping the terminal arbors of retinal axons in vivo. However, only cAMP modulations in PCs influence FAK phosphorylation on tyrosine 925. mean ± s.e.m.; individual data points are shown. * P<0.05, ** P<0.01; two-tailed Mann-Whitney test. Scale bar, 50 µm.

Overall, we have identified a PC specific cAMP signal that contributes to axon pathfinding. This local cAMP signal controls the density of PCs by influencing their assembly and disassembly rate in developing growth cones. Notably, we showed that PC-restricted cAMP signaling influences the phosphorylation of FAK on tyrosine 925 and that this phosphorylation event is required for ephrin-A5- evoked axon retraction. Finally, we demonstrated that lipid raft-specific cAMP signaling does not influence or overlap with the PC-restricted cAMP pathway controlling ephrin-A5-mediated axon guidance, suggesting that cAMP signals in different compartments regulates distinct cellular events that cooperate to fine tune the steering behavior of developing axons (**Figure 7c**).

## Discussion

### Diversity of subcellular-restricted cAMP signals

In addition to PCs, other subcellular domains containing local cAMP signaling have been identified in developing axons. A reduction of cAMP signaling restricted to lipid rafts is required for ephrin-A5-induced axonal retraction ^5^. In contrast, Slit1-induced growth cone collapse involves a cAMP-dependent pathway in a plasma membrane domain excluded from lipid raft ^6^. However, each domain-specific cAMP modulation is associated with a single axon pathfinding molecule and each guidance cue induces cAMP changes in a single cellular compartment. These subcellular domain-restricted cAMP signals also modulates different axonal behaviors ^6^. The identification of PCs as an additional compartment where cAMP regulates ephrin-A5-dependent axon repulsion highlights that several cAMP pathways cooperate to control the behavior of axons exposed to this guidance molecule. Interestingly, buffering cAMP in PCs and in lipid rafts in vivo do not lead to the same phenotypes, although both manipulations impact the development of the terminal arbor of retinal axons, a process that is dependent on the ephrin-A family ^25^. Altering lipid raft-specific cAMP pathways leads to an extended exuberance of axonal terminals in the SC, with long branches extending towards the caudal SC ^5^. In contrast, the terminal arbors of retinal axons in which cAMP is buffered in PCs are enlarged but do not exhibit long exuberant axonal branches (**Figure 7**). These distinct phenotypes further confirmed that cAMP pathways in these two cellular domains influence different molecular events involved in the ephrin-A-mediated control of axonal behaviors.

Since the regulation of cell adhesion modalities including FAK-dependent adhesion is not specific to ephrin-A5 but also occurs downstream of other axon guidance cues, including Netrin-1, Semaphorin 3A and ephrin-A1 ^10–13,23^, PC-restricted cAMP signals might be shared by several axon repellents and/or attractants, in contrast to lipid raft-restricted and lipid raft-excluded signaling ^6^. Other cellular compartments confining cAMP signaling have been identified in non-neuronal cells. These domains rely on a diversity of confining mechanisms including liquid-liquid phase separation ^26^ and local phosphodiesterase activity ^1^. Other cellular nanodomains are specific to individual receptors that associate their downstream effectors in their vicinity, thus forming distinct receptor-associated cAMP nanodomains ^2^. Similar domains might be found in developing axons thus extending further the range of cAMP compartments that modulate ephrin-A5-dependent axon repulsion. These mechanisms might also account for the confinement of cAMP signals in the vicinity of PCs.

### Molecular events controlled by local cAMP signaling

The diversity of the cellular domains containing specific cAMP signals might reflect the complexity of the cAMP-dependent pathways involved in axon pathfinding. The immediate cAMP- downstream events that regulate axon guidance include the main downstream effectors of this second messenger: PKA, members of the Epac family and likely cyclic nucleotide calcium channels ^20,21,27,28^. However, these proteins influence almost as many signaling pathways and cellular behaviors as cAMP. A diversity of more specific cellular events influencing axon pathfinding have been identified, without a link between these events and specific subcellular location of the cAMP source. This second messenger controls the dynamics of microtubules together with membrane remodeling in turning axons ^29^. cAMP-dependent vesicular transport controls the plasma membrane targeting of DCC, one of the Netrin-1 receptors, in axons exposed to this guidance molecule ^30^. In non-neuronal cells, the cAMP/PKA pathway directly regulates a non-canonical and ephrin-A-independent EphA2 pathway by increasing the phosphorylation of this ephrin-A receptor on a phosphorylation hotspot. The non-canonical signaling and the canonical ephrin-A binding-dependent pathways influence each other since an EphA2 receptor cannot induce both downstream pathways at the same time ^31^.

cAMP signaling is also associated with molecular determinants of cellular adhesion involved in axon pathfinding. Combined integrin activation and cAMP signaling cause growth cone collapse ^32^. Simultaneous activation of these two signaling pathways might occur downstream of Netrin-1 to influence axon steering ^33^. The control of this regulation might involve the cAMP-dependent kinase PKA since its activity influences the phosphorylation of FAK ^16^. It supports our findings describing a subcellular-restricted cAMP signal that controls FAK phosphorylation in the context of axon pathfinding. It is however unlikely that PKA directly induces the phosphorylation of the tyrosine 925 of FAK since the reduction in the PKA kinase activity increases the overall phosphorylation of FAK ^16^. A potential link between cAMP signaling and PC disassembly might rather be through calcium signaling. cAMP and calcium signaling often interact in neurons and non-neuronal cells ^34^. A substrate-dependent regulation of ryanodine receptor 3 by cAMP control the guidance behavior of axons from dorsal root ganglia neurons, thus linking cAMP to calcium signaling in an adhesion-dependent axon guidance context ^35^. In addition, calcium modulation has been associated with PC and focal adhesion disassembly. In non-neuronal cells, calcium transients trigger focal adhesion disassembly ^36–38^. In developing axons, this process might involve the calcium-dependent protease calpain that controls the balance between tyrosine kinase and phosphatase activities and influence the stability of PCs ^19,36,39^.

Other important regulators of PC dynamics might be involved in the regulation of PC remodeling downstream of ephrin-A5 and PC-restricted cAMP signaling. The cAMP-dependent kinase PKA is able to phosphorylate the three members of the Ena/VASP family, Mena, EVL and VASP ^40–42^. Ena/VASP proteins interact with the focal adhesion proteins zyxin and vinculin ^43,44^. They are required for integrin-dependent adhesion ^45^, and EVL largely contributes to cell-extracellular matrix adhesion and focal adhesion maturation during mechanically directed cell migration. In addition, the binding of EVL to the neuronal isoform of Src is abolished by PKA-dependent phosphorylation of EVL ^40^. Src is a kinase that is a direct interactor and regulator of FAK and is a downstream effector of the ephrin-A receptor EphA2 ^46^. Ena/VASP proteins might thus contribute to the regulation of PC dynamics by subcellular restricted cAMP signals, linking the latter to FAK phosphorylation.

### Impact of cAMP-dependent control of FAK phosphorylation at Y925

Previous investigations provided insights on how FAK phosphorylation on the tyrosine 925 might influence the density of PC. FAK phosphorylation on Y925 has been associated with axon repulsion ^13^, and with an increased stability of PCs in axons ^47^ and focal adhesions in non-neuronal cells ^48,49^. ^p925^FAK favors the disassembly of focal adhesions and slows down their turnover ^48^ by promoting Grb2 binding to FAK ^50,51^. This mechanism might be the downstream event controlled by PC-restricted cAMP signaling and explaining the impact of pax-cAMP Sponge on ephrin-A5-induced axon retraction in vitro and the refinement of RGC axon terminal arbors in vivo.

A prerequisite for Src to bind FAK and access Y925 is the activation of FAK, i.e. the autophosphorylation of FAK on Y397, followed by the exposure of the H1 helix of FAK, where Y925 is located ^52,53^. This mechanism provides an explanation for the reduced axon retraction of FAK when ^Y397F^FAK is expressed, although Y397 phosphorylation itself is not dependent on PC-restricted cAMP signaling.

In conclusion, we identify PCs as a subcellular domain containing a local cAMP signal that controls ephrin-A-dependent axon pathfinding through the regulation of cellular adhesion. This highlights that a diversity of cellular compartments confine signals of the same second messenger that control different signaling events in developing axons. Our findings demonstrate that subcellular compartmentation is crucial for cAMP to coordinate a diversity of axonal behaviors that wire the developing nervous system.

## Methods

### Animals

Pregnant C57BL6/J mice were purchased from Janvier Labs. All animal procedures were performed in accordance with institutional guidelines and approved by the local ethics committee (C2EA-05: Comité d’éthique en expérimentation animale Charles Darwin; protocol APAFIS#22331-2019100814127972v4). Animals were housed on a 12 h light/12 h dark cycle in temperature- and humidity-controlled environment (19–23 °C, 45–60%). Embryos from dated matings (developmental stage stated in each section describing individual experiments) were not sexed during the experiments and the female over male ratio is expected to be close to 1.

### Retinal explants

Retinas of E14.5 mouse embryos were electroporated using two poring pulses (square wave, 175 V, 5 ms duration, with 50 ms interval) followed by four transfer pulses (40 V, 50 ms and 950 ms interpulse) with a Nepa21 Super Electroporator (NepaGene). This electroporation procedure was used for the following plasmids: YFP, GFP, YFP-FAK, YFP-FAK Y925F, YFP-FAK Y861F, YFP-FAK Y397F, Paxillin-mRFP, Paxillin-GFP, Paxillin-cAMP Sponge-mCherry, Lyn-cAMP Sponge-mCherry and Lyn-mutated cAMP Sponge-mCherry (2 μg·μL^-1^ for single plasmid electroporation, 1 μg·μL^-1^ for each plasmid for co-electroporation). Retinas were dissected and kept 24 h in culture medium (DMEM-F12 without phenol red, supplemented with 1 mM glutamine (Sigma Aldrich), 1% penicillin/streptomycin (Sigma Aldrich), 0.001% BSA (Sigma Aldrich) and 0.07% glucose), in a humidified incubator at 37 °C and 5% CO2. The following day, the retinas were cut into 200 μm squares with a Tissue-Chopper (McIlwan) and explants were plated on 14 mm glass coverslips, 35 mm glass bottom dishes (Ibidi) or 4-well glass bottom slides (Ibidi), coated with 100 μg·mL^-1^ poly-lysine and 20 μg·mL^-1^ Laminin (Sigma Aldrich). Cells were cultured for 24 h in culture medium supplemented with 0.5% (w/v) methyl cellulose and B-27 (1/50, Lifetechnologies).

### Molecular biology

Lyn-cAMP Sponge-mCherry and its mutated variant were previously generated ^5^. All FAK and related phosphoresistant variants were obtain from Addgene (YFP- FAK: #50515, YFP-FAK- Y397F: #50516, YFP-FAK-Y861F: #50508, YFP-FAK-Y925F: #50509) and subcloned into a pcX backbone. Paxillin-mRFP and Paxillin-cAMPSponge-mCherry were generated using Paxillin-GFP backbone and replacing GFP sequence by mRFP and cAMP Sponge-mCherry ones.

### Collapse assay

Retinal explants were treated with 500 ng·mL^-1^ recombinant mouse EphrinA5 (R&D Systems) or PBS diluted in warm culture medium for 5 or 60 min before fixation with 4% (w/v) PFA in Sucrose 4% (w/v) for 20 min.

### Immunostaining following collapse assay

Retinal explants were permeabilized and blocked with 0.25% (v/v) Triton and 3% (w/v) BSA in PBS. All antibodies were diluted in PBS supplemented with 0.1% (v/v) Triton and 1% (w/v) BSA. For collapse assay, retinal explants were immunostained using a GFP antibody (1/1000, AvesLabs, Ref: GFP-1010) followed by a secondary antibody coupled to AlexaFluor 488 (1/500, Jackson Immunoresearch, Ref: 703-545-155) and a βIII-tubulin antibody (1/1000, Biolegend, Ref: 801202) followed by a secondary antibody coupled to AlexaFluor 647 (1/500, Jackson Immunoresearch, Ref: 715-605-150).

For TIRF imaging of fixed samples, retinal explants were immunostained using a GFP antibody (1/1000, AvesLabs, Ref: GFP-1010) followed by a secondary antibody coupled to AlexaFluor 488 (1/500, Jackson Immunoresearch, Ref: 703-545-155) and a RFP antibody (1/1000, Chromotek, Ref: 5f8-20) followed by a secondary antibody coupled to AlexaFluor 555 (1/1000, Invitrogen, Ref: A21434). The same samples were also immunolabeled with FAK/pFAK or Paxillin antibodies (FAK C-20, 1/200, Santa Cruz, Ref: sc-558. p-FAK(Y397), 1/500, Invitrogen, Ref: 44-624G. p-FAK(Y925), 1/200, Santa Cruz, Ref: SC11766. p-FAK(Y861), 1/500, Invitrogen, Ref: 700154. Paxillin, 1/500, BD Biosciences, Ref: 610051) followed by a secondary antibody coupled to AlexaFluor 647 (For FAK/pFAK: 1/500, Jackson Immunoresearch, Ref: 711-605-152. For Paxilin, the same as collapse assay).

### TIRF imaging and analysis

Images were acquired with an inverted Eclipse Ti2 (Nikon) coupled with a 100× oil-immersion objective, TIRF module (Gattaca system) and Metamorph software (MolecularDevices). Lasers were calibrated to 190 nm depth for every wavelength and images were taken using a Prime95B camera (Photometrics).

For live imaging, 2 μM HEPES at 37 °C was added to the retinal explant 20 min before imaging. Acquisitions were performed in a thermostatic chamber set at 37 °C. Images were taken every 5 sec during a 2 min time window. In order to match the timing done in collapse assay experiment, 3 series of acquisitions were made for each growth cone: 2 min before stimulation, around 5 min after stimulation (between 4 min 30 s and 6 min 30s) and around 10 min after stimulation (between 9 min 30 s and 11 min 30 s). Stimulation with 500 ng·mL^-1^ recombinant mouse ephrin-A5 (R&D Systems) or PBS diluted in warm culture medium was done manually. Analysis was performed using ImageJ and Manual tracking plugin. A Paxillin-GFP positive PC was considered as a static point, clearly distinguishable from the background (i.e. still visible when background is removed using ImageJ thresholding tool). Base on previously described PC dynamics ^10^ and ephrin-A5-induced retraction kinetics, only Paxillin-GFP positive points contact lasting at least 10 sec were considered for analysis. No cells were removed from the data set, except growth cone collapsing before any stimulation.

For fixed experiment, analysis was conducted using ImageJ ITCN plugin. GFP staining (allowing growth cone morphology visualization) was used as a mask for the plugin to define the counting area. The counting was done with the following parameters: Width = 6 pixels, Minimum distance = 3 pixels, and threshold = 1. The counting was systematically verified and corrected manually if needed using the same parameters.

### DIC imaging and analysis

Imaging was performed using an inverted DMI6000B Leica microscope, with a 40× oil-immersion objective. Recording medium was replaced with a laminar flux. Temperature was kept at 37°C during the whole recording. Retraction assays were performed by bathing the cells in a medium containing: 1 mM CaCl_2_, 0.3 μM MgCl_2_, 0.5 mM Na_2_HPO_4_, 0.5 mM NaH_2_PO_4_, 0.4 μM MgSO_4_, 4.25 mM KCl, 14 μM NaHCO_3_, 120 mM NaCl, 0.0004% CuSO_4_, 1.24 μM Fe (NO_3_)_3_, 1.5 μM FeSO_4_, 1.5 μM thymidine, 0.51 mM lipoic acid, 1.5 mM ZnSO_4_, 0.5 μM sodium pyruvate, 1 × MEM Amino Acids, 1 × non-essential amino acids, 25 mM Hepes, 0.5 mM putrescine, 0.01% BSA, 0.46% glucose, 1 mM glutamine, 2% penicillin streptomycin. Vitamin B12 and riboflavin were omitted because of their autofluorescence. DIC images were acquired using a CCD camera (ORCA-D2, Hamamatsu). Cells were imaged every 2 min for up to 90 min. Solution was perfused at a speed of 0.2 ml·min*^−^*^1^. Cells were bathed in control medium for 30 min to measure the growth rate and eliminate immobile growth cones from the analysis. A single fluorescent image was acquired before acquisition of identified electroporated axons. Only axons that grew faster than 60 μm·h^−1^ were included in the analysis. After 30 min, the culture medium was replaced by a medium containing ephrin-A5 (500 ng·μl*^−^*^1^), and axons were imaged for 1 h. Axon trajectories were tracked using a manual tracking plugin for ImageJ (NIH) and the retraction length was computed.

### In utero electroporation

Pregnant wild-type females were anesthetized with isofluorane. Using a glass micropipette (Dutscher), the left eyes of E14.5 embryos were injected with a mix of 2 DNA constructs: Pax-cAMP sponge (2.7 µg·µL^-1^) and GFP (1.1 µg·µL^-1^) or mRFP (2.7 µg·µL^-1^) and GFP (1.1 µg·µL^-1^). Retinas were electroporated with 5 pulses of 45 V for 50 ms every 950 ms (Nepagene electroporator). The positive electrode was placed on the injected eye and the negative electrode at the opposite side (CUY650P5, Sonidel) ^54,55^. Sub-cutaneous injections of flunixin-meglumine (4 mg·kg^-1^, Sigma) were applied for analgesia after the surgery. To increase the survival of the pups, a Swiss female mated one day earlier than the electroporated mice was used to nurse the pups. At P0, all but 3 Swiss pups were removed to stimulate nursing.

### Immunostaining following in utero electroporation

At P10, after anesthesia with pentobarbital (545 mg·kg^-1^), in utero electroporated pups were perfused transcardially with 4% paraformaldehyde (PFA) in 0.12 M phosphate buffer. Brains were postfixed overnight in 4% PFA and the whole superior colliculi (SC) were dissected, washed in PBS, permeabilized in 1% Triton for 30 minutes, blocked in 0.1% Triton 10% horse serum in PBS for 1h. The SC and retinas were incubated at 4°C for 3 days in antibodies raised against DsRed (1/800, Clonetech California; validated for similar assays in ^56^) and GFP (1/1000, Aves Lab; validated for similar assays in ^57^) diluted in the blocking solution. Finally, the SC were washed in PBS 3 times for 10 minutes each, incubated at room temperature for 2 hours in the secondary antibodies (Alexa 488 Donkey anti-Chicken, 1/200, Invitrogen; and CY3 Donkey anti-rabbit, 1/200, Jackson) diluted in the blocking solution and washed again in PBS 3 times for 10 minutes. SC were mounted in mowiol-Dabco.

### Whole mount SC imaging and analysis

Whole-mount SC were imaged using a Leica SP5 confocal and a 20X objective. Since the signal-to-noise ratio was higher in the GFP channel, electroporated axons were imaged using GFP staining. Individual arbors were identified and the position of the end of individual branches was manually pointed using ImageJ. The 3D convex hull and the total number of terminal branches was computed using ImageJ.

### Computational modeling

Each iteration of the simulation represented 5 seconds, matching the frequency of acquisition during imaging experiments. For each iteration, the number of new PCs was set to the measured appearance rate of PCs (normalized to the sham stimulated axons), with some variability introduced by a random number generator. The lifetime of each newly created PC was adjusted to the measured lifetime, thus determining the dissociation time of each PC and adjusting the dissociation rate to the rate of formation of new PCs, as measured (**Figure 3c**). During the 5 first minutes of the simulation (60 iterations), the rate of formation of PCs was stable and set to 1, to mimic the behavior of the growth cone before being exposed to ephrin-A5. From the 6^th^ to the 10^th^ minutes, it was set to a linear reduction reaching the measured rate 5 minutes after ephrin-A5 exposure. This rate was different in pax-RFP- and pax-cAMP Sponge-expressing axons (**Figure 3c**). From the 11^th^ to the 15^th^ minutes the appearance rate was set to follow a linear decrease reaching the measured frequency of PC formation measured after 10 minutes of ephrin-A5 exposure. After the 15^th^ minute, the rate of PC appearance was maintained stable until the end of the simulation. Using these parameters, 100 simulation rounds were averaged.

### Statistical analysis

No data were excluded from the analysis. No sample size calculation was performed. Sample size was considered sufficient after at least three independent experiments, leading to n ≥ 3 since several coverslips were often analyzed for the same experimental condition. Cultures were equivalent and not distinguishable before treatment, de facto randomizing the sample without the need of a formal randomization process. Image calculation and analysis were performed using ImageJ.

## Acknowledgements

We thank Dr Kenneth Yamada, Dr Rick Horwitz and Dr Aldebaran Hofer for the gift of the pax-GFP, phosphoresistant forms of FAK and cAMP Sponge plasmids. We are grateful to the members of the Nicol&Fassier lab for thoughtful discussion and helpful critical reading of the manuscript, and to the members of the animal and imaging facilities of Institut de la Vision. This work was supported, by grants from ANR (ANR-18-CE16-0017, ANR-22-CE16-0034), Fondation pour la Recherche Médicale (EQU202003010158) and Unadev (RGCMatch) to X.N. This work was performed in the frame of LABEX LIFESENSES (ANR-10-LABX-65) and of IHU FOReSIGHT (ANR-18-IAHU-0001) supported by French state funds managed by the Agence Nationale de la Recherche within the Investissements d’Avenir program. J.B. was supported by fellowships from the Fondation de France (00099274) and Fondation pour la Recherche Médicale (FDT202204014862). N.A.L.C. was supported by a fellowship from the Swiss National Science Foundation (P500PB_206674).

## Author Contributions Statement

Conceptualization, JB, XN; Methodology, JB, XN; Validation, JB, XN; Formal Analysis, JB, CM, XN; Investigation, JB, CM, AA, NALC, IK, CGB, FR, SC, XN; Writing – Original Draft, XN; Writing – Review & Editing, JB, CF, XN; Visualization, JB, XN; Supervision, XN; Project Administration, XN; Funding acquisition, JB, XN.

## Competing Interests Statement

The authors declare no competing interests.

## Notes

### Competing Interest Statement

The authors have declared no competing interest.

