## Supplemental Figures 1-3 for "Point contact-restricted cAMP signaling control ephrin-A5-induced axon repulsion"

### SUPPLEMENTARY FIGURES

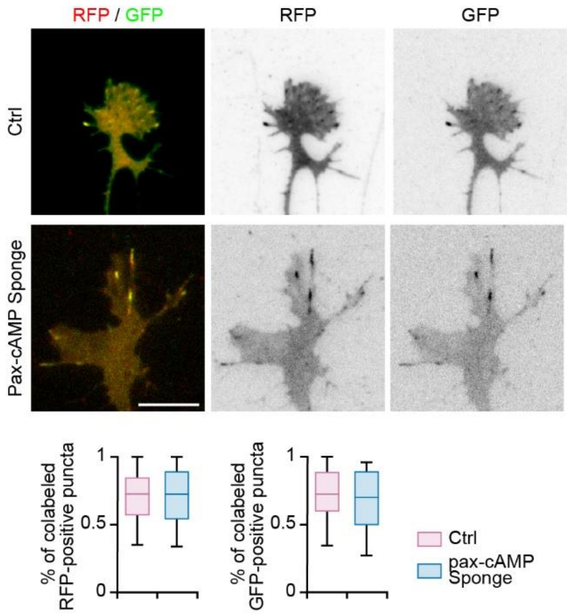


**Supplementary Figure 1. Subcellular targeting of pax-cAMP Sponge.**
The cAMP scavenger cAMP Sponge was fused to the full length paxillin (pax-cAMP Sponge). Pax-cAMP Sponge or its control pax-RFP were co-expressed with a previously validated marker of Focal adhesion or PC in living cells and axons. Pax-RFP, pax-GFP and pax-cAMP Sponge all label punctate structures in developing growth cones and the co-labeling behavior between pax-RFP and pax-GFP is similar to the behavior of the pax-cAMP Sponge/pax GFP pair. Box-and-whisker plot elements: median, upper and lower quartiles, $10^{th}$ and $90^{th}$ percentiles. Mann-Whitney test. Scale bar, 10 µm.

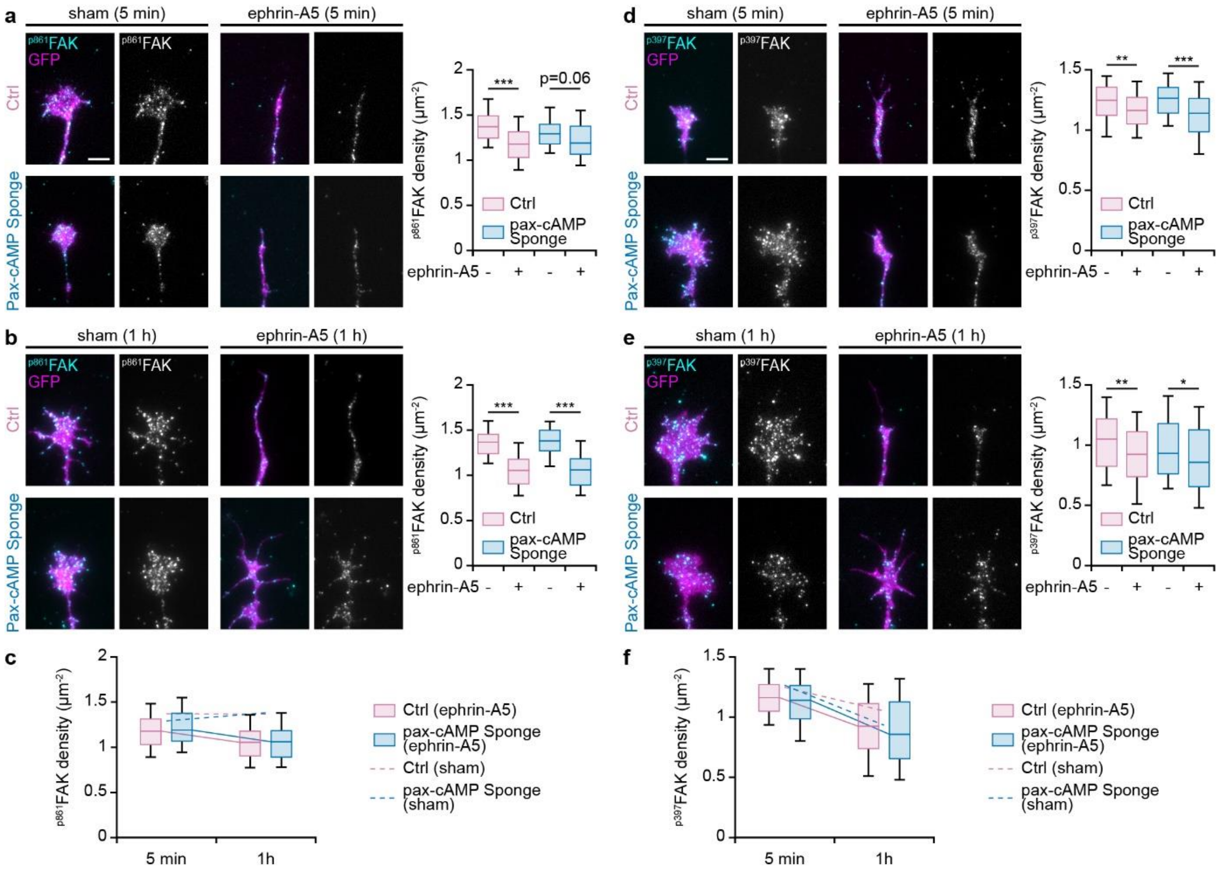


**Supplementary Figure 2. Pax-cAMP Sponge expression does not impact ephrin-A5-induced changes in FAK phosphorylation on tyrosines 861 and 397.**

Retinal axons expressing GFP together with either pax-RFP or pax-cAMP Sponge were exposed to ephrin-A5 or PBS as a sham control, immunolabeled for the phosphorylated tyrosines 861 or 397 of FAK and imaged using TIRF microscopy. **(a)** Ephrin-A5 induces a reduction in the density of $^{p861}$FAK 5 min after the stimulation, in pax-RFP-expressing growth cones. A similar density of $^{p861}$FAK was detected in pax-cAMP Sponge-expressing growth cones. **(b)** After long term exposure to ephrin-A5 (1 h), the density of $^{p861}$FAK-positive PCs was reduced compared to PBS treated axons and no difference was detected between pax-mRFP- and pax-cAMP Sponge-positive axons. **(c)** Summary of the changes in $^{p861}$FAK-positive PCs over time. **(d)** Ephrin-A5 induces a reduction in the density of $^{p397}$FAK 5 min after the stimulation, in pax-RFP-expressing growth cones. A similar density of $^{p397}$FAK was detected in pax-cAMP Sponge-expressing growth cones. **(e)** After long term exposure to ephrin-A5 (1 h), the density of $^{p397}$FAK-positive PCs was reduced compared to sham-stimulated axons and no difference was detected between pax-mRFP- and pax-cAMP Sponge-positive axons. **(f)** Summary of the changes in $^{p397}$FAK-positive PCs over time. Box-and-whisker plot elements: median, upper and lower quartiles, 10$^{th}$ and 90$^{th}$ percentiles. * $P<0.05$; ** $P<0.01$; *** $P<0.001$; Kruskal-Wallis test followed by Dunn's post-hoc test. Scale bar, 5 µm.

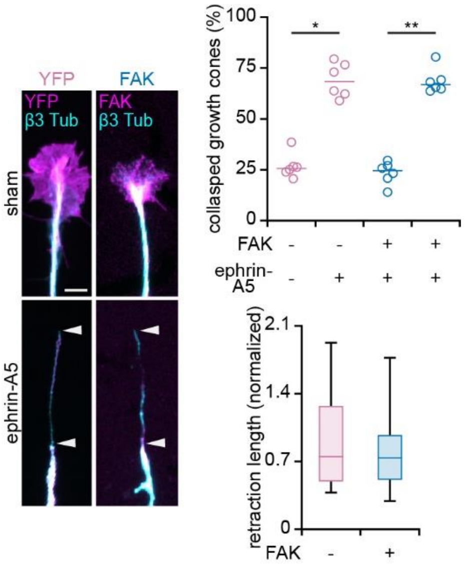


**Supplementary Figure 3. Overexpressing FAK does not affect ephrin-A5-induced growth cone collapse and axon retraction**

Axons expressing YFP or FAK were exposed to ephrin-A5 for 1 hour. YFP- and FAK-overexpressing growth exhibited similar percentage of collapse, and the retraction process of collapsed had similar length, indicating that the retraction of these axons is not affected by FAK overexpression. Top graph; median and individual values; Kruskal-Wallis test followed by Dunn's post-hoc test. Bottom graph; Box-and-whisker plot elements: median, upper and lower quartiles, 10th and 90th percentiles; Mann-Withney test. * $P<0.05$; ** $P<0.01$; Scale bar, 5 µm.
